# Dead or gone? Bayesian inference on mortality for the dispersing sex

**DOI:** 10.1101/031161

**Authors:** Julia A. Barthold, Craig Packer, Andrew J. Loveridge, David W. Macdonald, Fernando Colchero

## Abstract

1. Estimates of age-specific mortality are regularly used in ecology, evolution, and conservation research. However, existing methods to estimate mortality from re-sighting records of marked individuals fail at estimating mortality of males for species with male natal dispersal due to the uncertainty surrounding disappearances of adult males from study populations.
2. Here, we develop a mortality model that imputes dispersal state (i.e., died or left) for uncertain male records as a latent state jointly with the coefficients of a parametric mortality model in a Bayesian hierarchical framework. To check the performance of our model, we first conduct a simulation study. We then apply our model to a long-term data set for African lions. Using these data, we further scrutinise the mortality estimates derived from our model by incrementally reducing the level of uncertainty in the male records. We achieve this by taking advantage of an expert’s intuition on the likely fate of each uncertain male record.
3. We find that our new model produces accurate mortality parameters for simulated data of varying sample sizes and proportions of uncertain male records. From the empirical study we learned that our model provides similar mortality estimates for different levels of uncertainty in male records. However, a sensitivity of the mortality estimates to varying uncertainty is, as can be expected, detectable.
4. We conclude that our model provides a solution to the challenge of estimating male mortality in species with data-deficiency for males due to natal dispersal. Given the utility of sex-specific mortality estimates in biological and conservation research and the virtual ubiquity of sex-biased dispersal, our model will be useful to a wide variety of applications.

## Introduction

Mortality estimates of both sexes for wild animal populations are fundamental for testing hypotheses derived from ecological and evolutionary theory, and for predicting population size and structure for population management purposes. However, estimating mortality of at least one of the sexes is commonly hindered by incomplete data on dispersing individuals. In many large vertebrate species, males leave their natal place or social group around the age of maturity. If dispersing males leave the areas monitored by field studies that collect re-sighting data on marked individuals, these migrating males impede the quality of gathered data. Following dispersing males using telemetry or GPS technology is cost and labour intensive, and therefore dispersing males are usually lost for data collection.

The possibility that missing males may have dispersed increases the uncertainty of the fate of all males that are no longer detected. Missing females are likely dead, even if their bodies are not found, since they do not disperse. Missing males, however, that were old enough for dispersal may have died or dispersed. This uncertainty in the male records prevents the estimation of male mortality using existing methods.

Models to infer survival using capture-mark-recapture/re-sighting (CMRR) data derived from the Cormack-Jolly-Seber framework (CJS; after Cormack, 1964; Jolly, 1965; Seber, 1965) can accommodate both uncensored and right-censored records (i.e., individuals known to be alive after the last observation). These approaches exploit the fact that each type of record contributes different information (White & Burnham, 1999). Extensions to the initial models have been developed that accommodate species-specific life histories and data issues arising from the movement of the individuals in relation to the spatial and temporal distribution of the marking and re-sighting effort. Accordingly, these models incorporate for example incomplete and heterogeneous re-sighting probabilities, multiple states, and multiple locations (e.g., Arnason, 1973; Schwarz, Schweigert & Arnason, 1993; Lebreton & Pradel, 2002; Mackenzie *et al*., 2009; Cubaynes *et al*., 2010; Lagrange *et al*., 2014; Ergon & Gardner, 2014). Furthermore, Schaub & Royle (2014) have recently developed a spatially explicit Cormack-Jolly-Seber approach that jointly models mortality and dispersal using movement data for species in which dispersal can be described as a random walk. None of these approaches can accommodate the uncertain male records that are typical for re-sighting data of species with male natal dispersal, where dispersal probabilities vary with age.

In order to address issues with missing records in CMRR data, Bayesian approaches have been developed that estimate survival probabilities and transition probabilities between states and locations while augmenting data (Dupuis, 1995, 2002; King & Brooks, 2002). Among these approaches, very flexible are those that estimate latent (unknown) states jointly with all other model parameters in a hierarchical framework using Markov Chain Monte Carlo (MCMC) algorithms (Clark *et al*., 2005; Colchero & Clark, 2012; Colchero, Jones & Rebke, 2012). Since latent states can be both finite sets of discrete states (e.g., locations or stages) or continuous variables (e.g., date of birth or death), this framework is suitable for developing a survival model that treats dispersal as a latent state, and can therefore accommodate uncertain male records.

Among mammals, males commonly form a data-deficient subpopulation. Male dispersal hinders inference on male mortality, and obscured paternities hinder inference on male lifetime reproduction. The lack of data on males prevents studies that would use measures of male fitness to test evolutionary theory or would use male demographic rates to address the role of males in population dynamics. Yet, in many mammal species, males are under stronger sexual selection than females, which makes them particularly interesting to study from an evolutionary perspective (Andersson, 1994). Furthermore, males have been found to influence the dynamics of populations through mechanisms such as sexual harassment and limiting female fecundity (Milner-Gulland *et al*., 2003; Le Galliard *et al*., 2005), and several more mechanisms are predicted by ecological theory (Mysterud, Coulson & Stenseth, 2002; Rankin & Kokko, 2007). As long as intensified data collection is out of reach, the development of statistical methods to infer male life history parameters from incomplete data is a timely endeavour.

Here, we present a model that can estimate age-specific mortality for both sexes in species with uncertain male records due to male dispersal. The model fits a parametric mortality model as a function of age and sex in a Bayesian hierarchical framework jointly with the estimation of the distribution of ages at dispersal, treating potential dispersal of males with uncertain records as a latent state. Using simulated data, we first validated the model. We then applied the model to estimate age-specific mortality of both sexes for Serengeti lions (*Panthera leo*) in Tanzania. Since this particular data set contains the expert opinion from the head of the study (C. Packer, unpublished data) on whether a missing male is likely to have dispersed or died, we used this information to gain further insights into the workings of our method.

## Methods

We focus on species in which males disperse only once at around the age of maturity (‘natal dispersal’). To isolate the effect of uncertainty in male records on mortality estimates from other effects, we focus on data that meet the following assumptions. We assume that individuals are re-sighted with certainty if they are alive and in the study area and that individuals are only observed at one location. We further assume that mortality in- and outside of the study area is equal, and that individuals born outside of the study area disperse into the study area with equal probabilities as individuals born in the study area disperse out of it. We also assume that ages of individuals whose birth was not observed (left-truncated records) can be estimated with sufficient certainty by a trained observer to allow us to not include time of birth as a latent state in the model and to model ages at death as a continuous variable. However, since the data available to us for the empirical application contained individuals that died before sexing was possible, we did construct the model to accommodate this type of record, treating the sex of unsexed individuals as another latent state. Finally, we further make one assumption that we know is not met for data from wild animal populations, and that is that mortality only depends on age and sex and not on any other covariates. However, this assumption allows us to develop a model to estimate baseline mortality for pooled data, that can later on be easily extended to incorporate other covariates, data quality permitting.

## Life History Data

### Data structure

The life history data used to estimate age- and sex-specific mortality included records for native-borns and immigrants. Native-borns were born in the study population, defined as all individually recognisable and constantly monitored individuals. Immigrants entered the study population some time after their birth either due to migration, or by being alive at the start of the study (Figure 1). The recorded types of departure from the population included death, censoring due to being alive at the end of the study, or uncertain fate (death or censoring through dispersal). Uncertain fates through dispersal were only caused by dispersals from the study population to an external population, and not by dispersals within the study population. Here, we refer to this out-migration from the study population when we use the term ‘dispersal’

**Figure 1:**
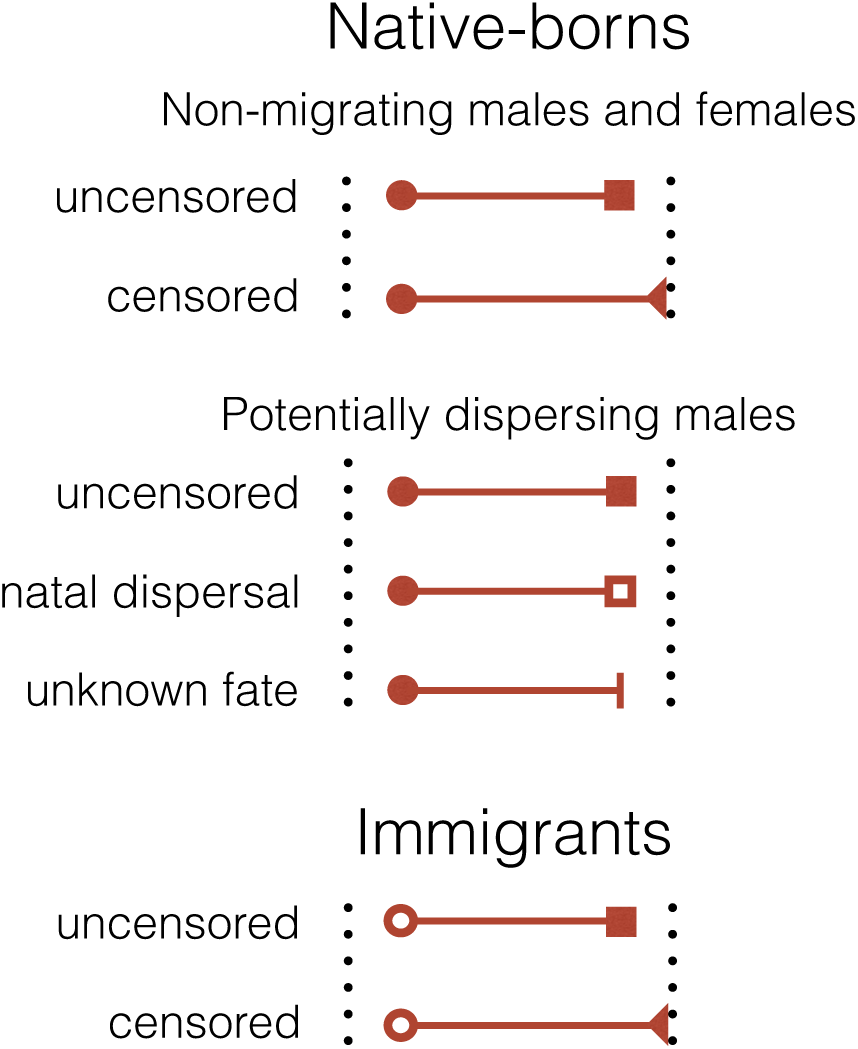
Example of types of records in the lion dataset. Circles represent times of entry 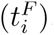, depending on its sex and dispersal statused circles corresponds to known times of birth and open circles are entries after birth (i.e., immigration or birth before the study started). Squares are departure times 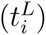 where filled squares are known times of death and open squares are dispersal. Filled triangles indicate individuals known to be alive at the end of the study and vertical bars indicate that the type of departure from the study population is uncertain (i.e., either death or dispersal).

### Simulated data

To validate the performance of our model, we used known mortality parameters to simulate data of the described structure and checked whether our model accurately retrieved these parameters. To simulate the data, we first randomly assigned a sex for an initial number of individuals by drawing from a binomial distribution, assuming an equal probability of being born male or female. We then randomly drew ages at death (*x_i_*) for each individual i by inverse sampling from a Siler CDF (see equations (2b) and (3)) with parameters ***θ****_f_* = {−1.4, 0.65,0.07,−3.8, 0.2} for females and *θ_m_* = {−1.2, 0.7, 0.16,−3.5, 0.23} for males. The subscripts *f* and *m* denote females and males, respectively. We then randomly drew ages at dispersal for all males by inverse sampling from a gamma CDF with parameters γ = {10, 3} and adding the minimum age of dispersal *α* = 1.75). We assigned every individual a last seen age 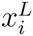 depending on its sex and dispersal status. For females and for those males whose ages at death were simulated to be younger than their ages at dispersal (i.e., they died before they could disperse), the last seen ages were the ages at death. For the other males, who were simulated to have died after dispersal, the last seen ages were set to be the ages at dispersal. Finally, to add immigrants to the data, we simulated the same number of males being born in the external population. For these males, as before, we randomly drew ages at deaths and ages at dispersal, and if they were simulated to have dispersed before death, we added them to the data as immigrants with their ages at death recorded as last seen ages and their ages at dispersal recorded as first seen ages 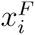.

We simulated data sets of two different initial numbers of native-borns (small sample size N = 500 and large sample size N = 2000). Within each sample size, we also produced further data sets where the sexes of all individuals were known, and data sets where we randomly assigned, with a probability of 0.3, the state of ‘unknown sex’ to all individuals that died at < 1 year of age. Finally, we simulated data that varied in the proportion of observed or ‘known’ deaths among individuals that were no longer re-sighted. We used three settings: 1, 5, and 10 % known deaths. In total, we thus simulated 12 data sets.

### Serengeti population

The study population occupied a 2000 km^2^ region of Serengeti National Park, Tanzania, that lies at the heart of the Serengeti-Mara ecosystem. The study site is characterised by seasonal rainfall and a southeast to northwest gradient in vegetation from short to tall grassland to open woodlands (Packer, 2005; Mosser *et al*., 2009). We analysed demographic data collected between 1966 and 2013. Observations were opportunistic between 1966 and 1984, and most animals were sighted 1-3 times per month. Study prides have been monitored with radio telemetry since 1984, allowing each animal to be observed 2-6 times per month. All individuals are identified from natural markings (Packer *et al*., 1991), and birth dates of cubs born in the study area are deduced from lactation stains on the mothers. A large number of nomadic males enter the area, and a small proportion become resident in one or more of the resident prides. Our analyses exclude all nomadic males that never became residents in the study area (N = 548). Individuals with unknown dates of birth were assigned an estimated age by a trained observer, using age indicators (e.g., relative body size, nose coloration, and eruption and wear of teeth) (Smuts, Anderson & Austin, 1978; Whitman *et al*., 2004). The data set contained a large number of individuals of unknown sex. Since the vast majority of these unsexed individuals died within the first weeks after birth, we excluded all individuals with last seen ages younger than 0.25 years of age. The final data set contained observations on 1341 females, 1263 native-born males, 316 immigrants, and 269 unsexed native-born individuals. The proportion of females among all native-born individuals (excluding immigrants), assuming a sex ratio of 1 to 1 among individuals that died before their sex could be determined, was 0.51.

## Mortality analysis

### Model variables and functions

We used a Bayesian approach that allowed us to estimate not only parameters for mortality and dispersal but also the different latent states such as dispersal state and sex. We used a parametric model to infer age-specific mortality. With *X* being a random variable for age at death, and x any age, the model required defining the mortality function or hazard rate as

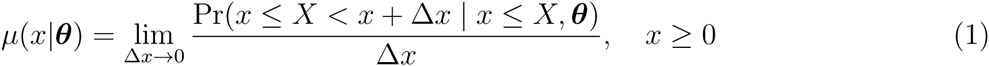
where ***θ*** is a vector of mortality parameters to be estimated. The estimated mortality can be used to calculate the probability to survive from birth to age x, or survivor function,

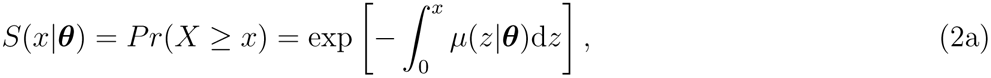
the probability that death occurs before age x, or the cumulative density function (CDF),

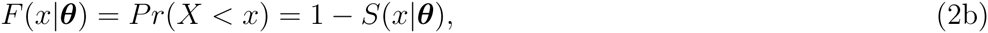
and the probability density function (PDF) for age at death

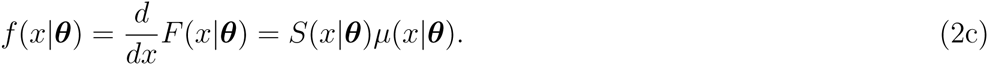

To capture the bathtub-shaped mortality rates typical of large mammals, we used the Siler model (Siler, 1979) in the form

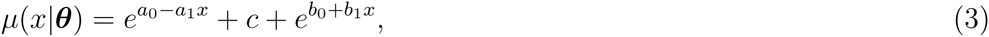
where ***θ*^⏉^** = [*a*_0_, *a*_1_, *c, b*_0_*, b*_1_], with 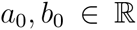 and *a*_1_, c, *b*_1_ > 0. The Siler model is a competing risk model constituted by three additive mortality hazards. The parameters capture different aspects of the shape of the age trajectory with *a*_0_ being the initial level of mortality rates and a_1_ governing the decrease in mortality over infant and juvenile ages. The *c* parameter scales mortality rates up or down and is usually interpreted as reflecting age-independent causes of mortality. This parameter is also dominant in capturing mortality in early adult ages when infant mortality has declined and senescence mortality not yet risen. The *b*_0_ parameter represents the initial mortality of the age-dependent increase of mortality and *b*_1_ determines the rate of this increase (Siler, 1979).

To model the ages at dispersal, we defined the random variable *Y* for age at dispersal, where the age at natal dispersal was *Y ~ G_Y_*(*y*) for ages *y* > 0, with *G_Y_*(*y*) being the Gamma distribution function with parameter vector γ^⏉^ = [γ_1_, γ_2_]. This distribution yielded the probability density function (PDF) of age at natal dispersal given by

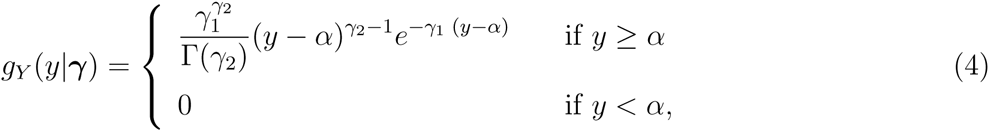
where *α* is the minimum age at natal dispersal and γ_1_, γ_2_ > 0. A summary of all the functions, parameters, indicators, and variables is provided in Table 1.

**Table 1:** Description of random variables, observed variables, and indicators.

| ***Modelled random variables*** |  |
| --- | --- |
| $X$ | Random variable for age at death, where $x$ is any age element |
| $Y$ | Random variable for age at natal dispersal with elements $y$ |
| $D$ | Binary random variable for disperser or non-disperser |
| $S$ | Binary random variable for sex |
| ***Observed variables and indicators*** |  |
| $\mathbf{t}^F$ | Vector of times of first detection |
| $\mathbf{t}^L$ | Vector of times of last detection |
| $\mathbf{b}$ | Vector of times of birth |
| $\mathbf{x}^F$ | Vector of ages at first detection ($x_i^F = t_i^F - b_i$) |
| $\mathbf{x}^L$ | Vector of ages at last detection ($x_i^L = t_i^L - b_i$) |
| $\mathbf{m}$ | Indicator vector for immigrants ($m_i = 1$ if immigrant) |
| ***Updated indicators*** |  |
| $\mathbf{d}$ | Indicator vector for dispersers ($d_i = 1$ if disperser and $d_i = 0$ otherwise) |
| $\mathbf{s}$ | Indicator vector for sex ($s_i = 1$ if female and $s_i = 0$ otherwise) |
| ***Parameters*** |  |
| $\boldsymbol{\theta}$ | Vector of mortality parameters |
| $\boldsymbol{\gamma}$ | Vector of natal dispersal parameters |
| ***Functions*** |  |
| *Mortality* |  |
| $\mu(x \mid \boldsymbol{\theta})$ | Mortality (Siler model) |
| $S(x \mid \boldsymbol{\theta})$ | Survival |
| $F(x \mid \boldsymbol{\theta})$ | CDF for age at death ($F(x) = 1 - S(x)$) |
| $f(x \mid \boldsymbol{\theta})$ | PDF for age at death |
| *Dispersal* |  |
| $g(y \mid \boldsymbol{\gamma})$ | PDF for age at natal dispersal (gamma distribution) |
| $G(y \mid \boldsymbol{\gamma})$ | CDF for age at natal dispersal |

### Likelihood and posterior

To construct the mortality likelihood, we assigned a different probability to each type of record in Figure 1. The likelihood for females and non-migrating nativeborn males was

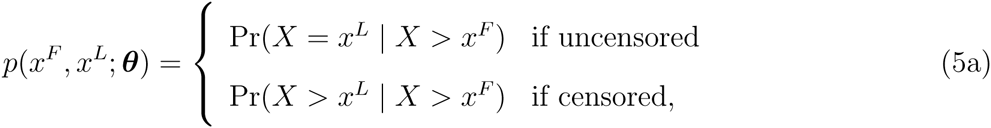
where *x^L^* corresponds to the age at last detection and *x^F^* is the age at first detection (i.e., *x^F^* = 0 for individuals born in the study area and *x^F^* > 0 for immigrants or individuals born before the study started), while for dispersers the likelihood was constructed as

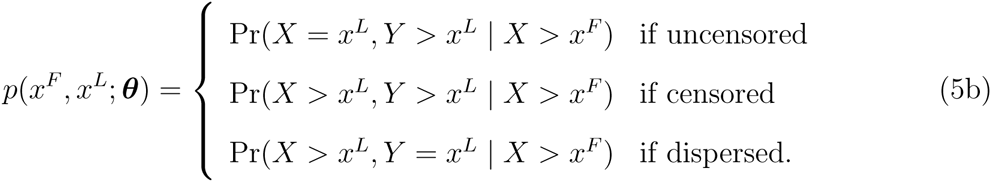

We defined dispersal state as a random variable *D* which assigned 1 if an individual *i*, born at *b_i_* and last detected at 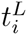, dispersed in its last detection age, 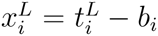, and 0 if otherwise. It was treated as a latent variable that needed to be estimated for males with uncertain fate. The censored and uncensored probabilities for dispersers were used to determine how likely it was for a potential disperser (i.e., departure type of uncertain fate and age older than minimum age at dispersal) to have dispersed (e.g., last expression in equation 5b) or died at the age of last detection *x_L_* (e.g., first expression in equation 5b). Furthermore, we also defined a binary variable S for the sex of the individual.

With this, we could construct the full Bayesian model as

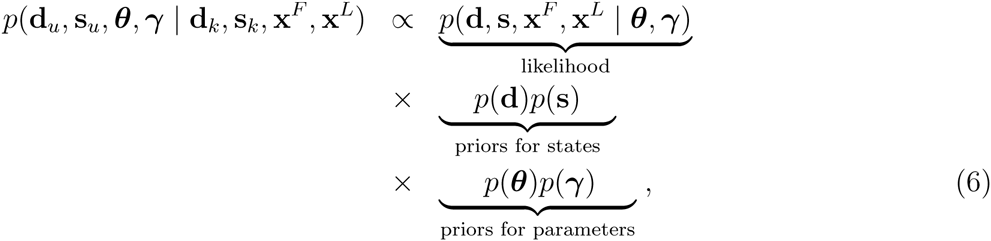
where d was the vector of dispersal states and s was the indicator vector for sex (*s_i_* = 1 if female and *s_i_* = 0 if male). Each of these vectors had two subsets represented by the subscripts *u* for unknown and *k* for known.

### MCMC and conditional posteriors

We used a Markov Chain Monte Carlo (MCMC) algorithm to fit the model in equation 6. For all implementations, we ran four parallel MCMC sequences with different randomly drawn starting values and set the number of iterations to 15,000 steps with a burn-in of 5,000 initial steps and a thinning factor of 20. We used a hierarchical framework that only needed the conditionals for posterior simulation by Metropolis-within-Gibbs sampling (Gelfand & Smith, 1990; Clark, 2007). This means that, for this particular case, the algorithm divided the posterior for the joint distribution of unknowns into four sections: (a) estimation of mortality parameters, (b) estimation of dispersal parameters, (c) estimation of unknown dispersal state, and (d) estimation of unknown sexes. Here we present each section, specifying the conditional posterior and the acceptance probability for the Gibbs Sampler algorithm.

#### Section a: Posterior for mortality parameters

The conditional posterior to estimate the mortality parameters **θ** required only the ages at last detection 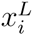 and the dispersal states *d_i_*. The posterior for a given individual *i* was

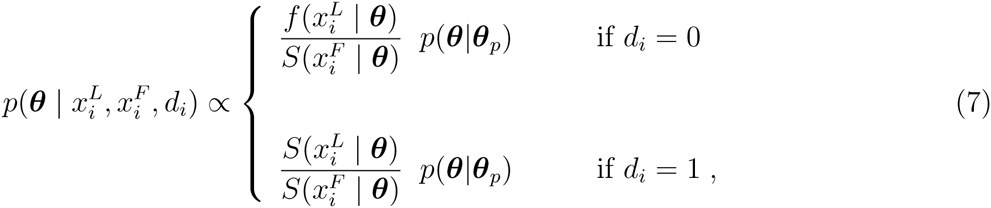
where ***θ****_ρ_* was a vector of prior hyper-parameters and 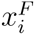 the age at first detection. If the individual was a native-born, then 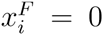 and the denominator in both expressions was equal to 1. At every iteration and for a given parameter *θ* ∈ ***θ*** with conditional posterior *p*(*θ* |…), the algorithm proposes a new parameter value for each element of *θ*′ and accepts it with acceptance probability

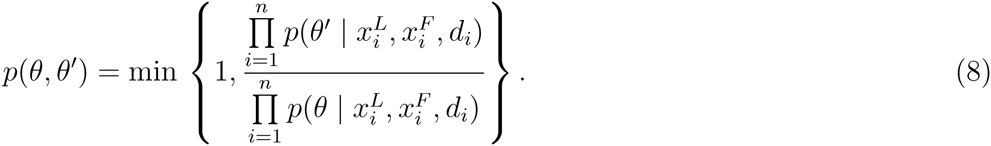

#### Section b: Posterior for dispersal parameters

The conditional posterior to estimate the parameters γ for the distribution of ages at first dispersal for a given individual *i* was

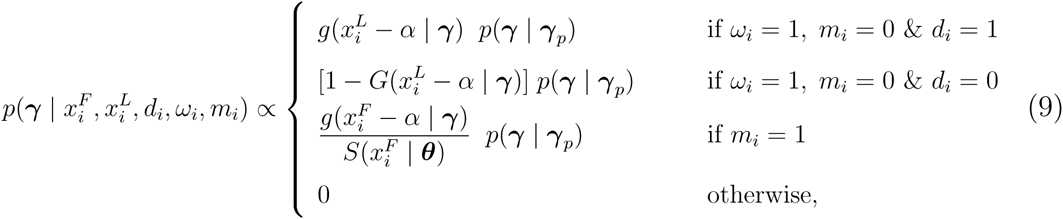
where γ*_p_* was a vector of prior hyper-parameters for γ, *ω_i_* was an indicator that assigns 1 if an individual was a potential disperser (i.e., if it belonged to the dispersing sex and disappeared at an age older than the minimum age at dispersal *α*), and *m_i_* was an indicator for immigrants. We set the minimum age at dispersal to *α* = 1.75 years for the simulated data and *α* = 1·5 for the Serengeti data. The age α corresponded to the earliest age at which immigrants could be detected and potential dispersers could be last seen. For a parameter γ ∈ γ with conditional posterior density *p*(γ |…) The acceptance probability for a proposed parameter of γ′ was

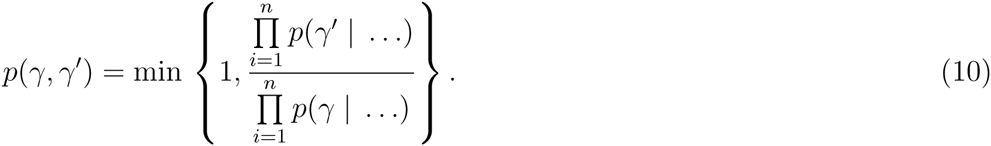

#### Section c: Posterior for dispersal states

Dispersal state was evaluated for individuals that were potential dispersers (i.e., *ω_i_* = 1) and estimated the joint probabilities

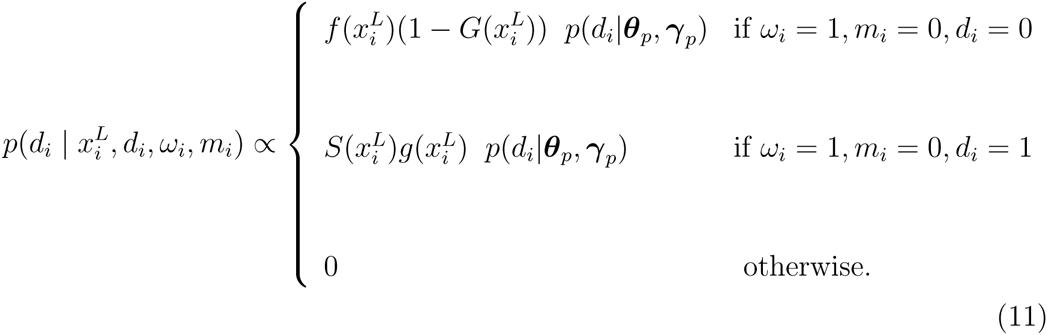

The first terms on the right hand side of equation 11 correspond to the likelihood function as defined in equations 5, while the second terms are the priors for dispersal state. For this section the acceptance probability for the sampling given the last seen ages, the dispersal states, the potential disperser states, and the immigration states was

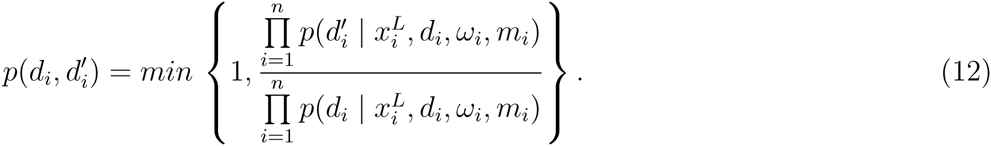

#### Section d: Posterior for unknown sexes

Some individuals disappeared before the minimum age at dispersal without their sex being determined. The conditional posterior for the latent state of sex was

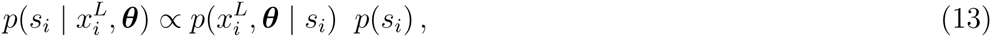
where the second term on the right-hand side is a prior for sex based on sex ratio at birth, or if the analysis was conditioned on survival to age *x*, based on the sex ratio at age *x*.

The indicator for potential dispersers *ω_i_* (see section *c*) was updated in each iteration. Individuals of undetermined sex and last seen ages older than the minimum age at dispersal were assigned 1 if imputed to be male and 0 if imputed to be female. The acceptance probability given the last seen ages and the mortality parameters was

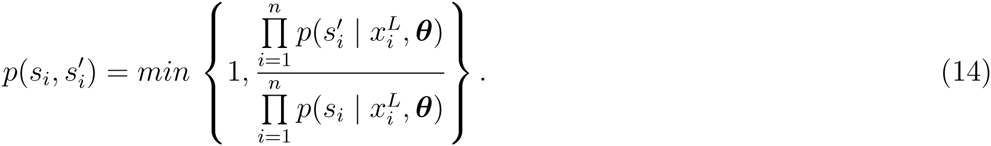

### Mortality and dispersal priors

We set the Siler parameters for the prior for females to 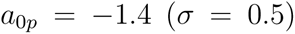, 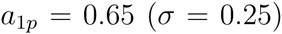, 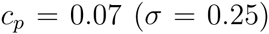, 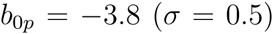 and 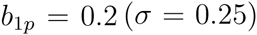, and for males to 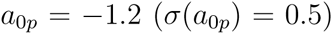, 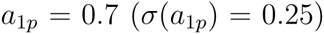, 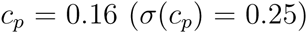, 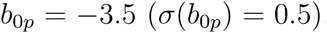 and 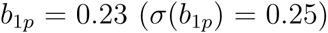. For dispersal, the Gamma parameters (shape and scale) for the prior were set to γ*_p_* = {8, 2} with σ(γ*_p_*) = {2,1}. Both the mortality and dispersal priors were uninformative.

### Model application and posterior analysis

We fitted the model with sex as a fixed covariate to the Serengeti data. In order to gain deeper insights into the performance of our model, we further exploited a unique source of information that is contained in this data set. A Serengeti lion expert used the circumstances accompanying the disappearances of males to deduce whether the individuals may have dispersed. For example, since young males often leave their natal prides with brothers, a simultaneous disappearance of brothers hints that this is likely to be a dispersal event. We fitted the model with three different settings. First, all males with uncertain fates and last seen ages older than minimum age at dispersal were assigned the state of ‘potential dispersers’ and entered in the model as described in ‘Section c’ above (Model A). Second, all males that were indicated to may have dispersed were entered as ‘known dispersers’ (see equation 5b) (Model B). And third, all males that were indicated to may have dispersed were entered as ‘potential dispersers’ (Model C).

To avoid problems arising from the large number of unsexed individuals that died within the first weeks after birth, we fitted the model from the start age of 0.25 years. We predicted mortality rates for each sex using the parameter estimates of every step of the MCMC after burn-in and thinning and used these predictions to calculate mean and credible intervals of mortality rates. To compare the three models to each other, we computed the life expectancy at the model start age and the Kullback-Leibler (KL) divergences of the mortality parameter posterior densities (Kullback & Leibler, 1951; McCulloch, 1989; Burnham & Anderson, 2001) (see Methods S1 for details on the calculation and the interpretation of KL values).

## Results

### Simulation study

We used a simulation study to validate our model. For all 12 simulations, the mortality rates used to simulate the data lay within the 95 % credible intervals of the estimated mortality for both sexes (Figure 2). Of all the introduced variations in data quality (sample size, unsexed individuals, proportion ‘known’ deaths), the only one with a marked effect on the performance of the model was varying the sample size. As could be expected, smaller sample sizes resulted in wider credible intervals particularly for males and for older ages of females. Due to the wider confidence bands for smaller sample sizes, the respective estimated mortality rates could appear to be less variable over the life span than the mortality rates used to simulate the data. This manifested as a less-pronounced U-shape of the estimated mortality rates when compared to the ‘real’ mortality rates (e.g., second panel in second row of Figure 2). The proportion of unsexed individuals dying at < 1 year of age, and the proportion of known deaths among disappearances did not discernibly affect the retrieval of the mortality parameters.

**Figure 2:**
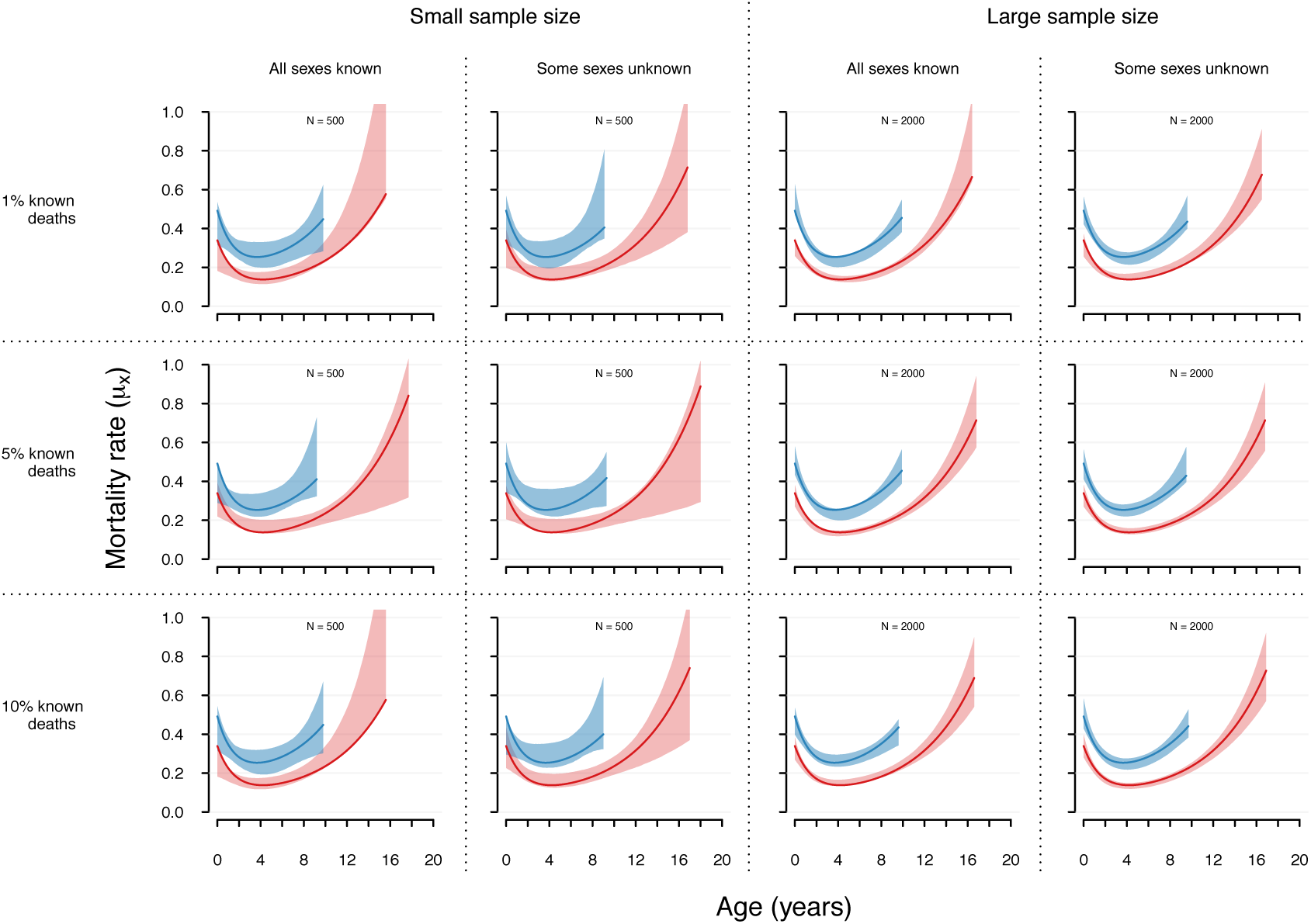
Predicted mortality rates for males (blue polygons) and females (pink polygons) compared to the mortality rates used to simulate the data (solid lines). Polygons represent 95 % credible intervals of age-specific mortality rates. Mortality rates are plotted until the ages when 95 % of a synthetic same-sex cohort would be dead. Results are given for 12 simulations varying the size of the native-born population (N = 500 or N = 2000), the proportion of known deaths among last seen ages (1%, 5%, or 10%), and whether the sex of 30% of individuals dying younger than 1 year of age remained undetermined or not.

### Application

The empirical models for Serengeti lions converged for all estimated parameters (Figure 3, see also Figure S1 for traces). Overall mortality of both sexes was U-shaped with high initial cub mortality, low mortality of prime-aged adults, and an age-dependent increase in mortality during the older ages (Figure 4). Mortality of males was higher than mortality of females across all ages (Figure 4), except for very young ages, up until one year, during which confidence bands of male and female mortality overlapped. However, this may be due to the large proportion of unsexed individuals at these ages (see data description) and the imputation of sex as a latent state for these individuals, which introduced uncertainty. Due to the higher male mortality rates across most ages, female life expectancy (4.7 years at model start age) exceeded that of males by approximately 2 years.

**Figure 3:**
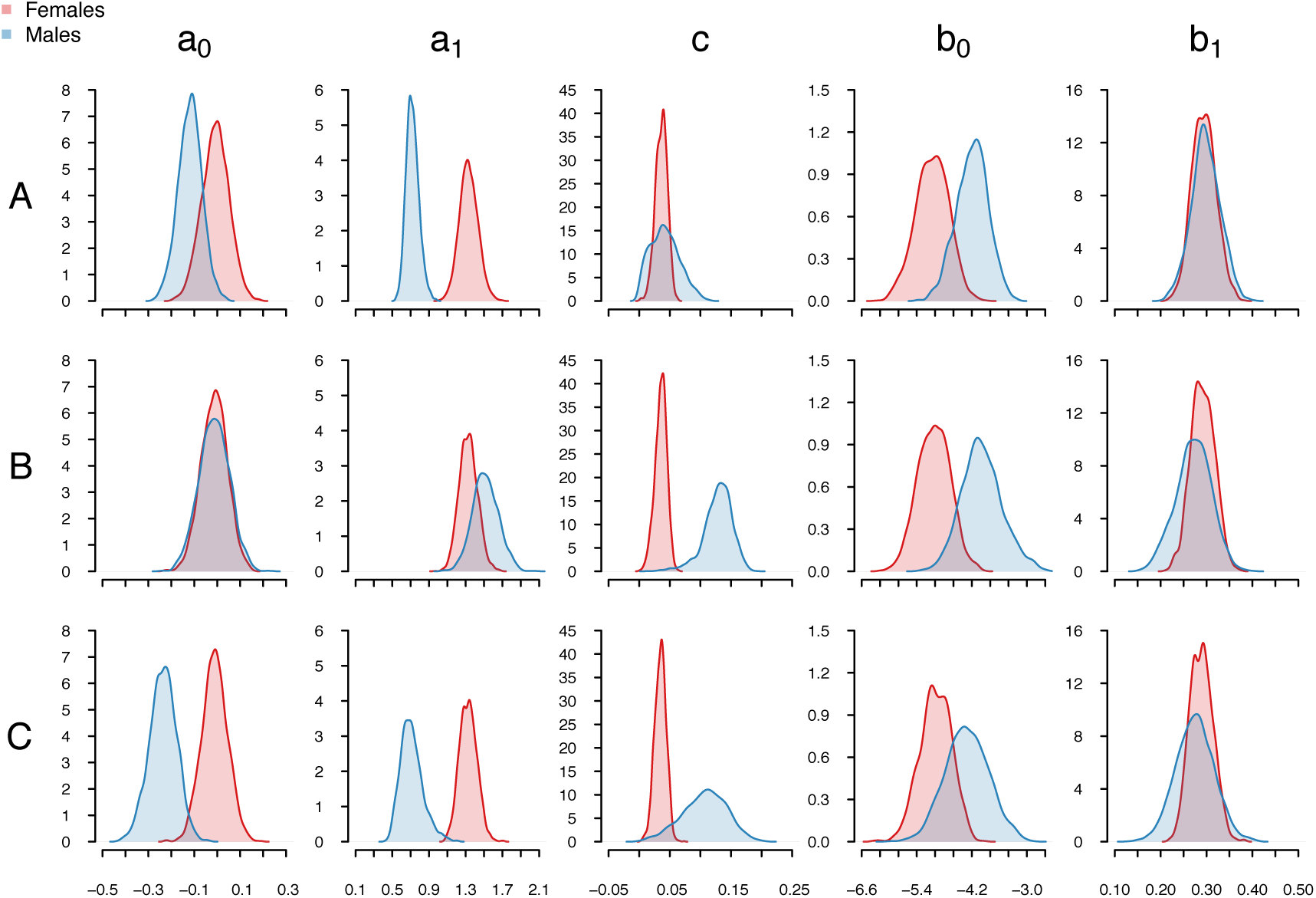
Posterior distributions of Siler parameter estimates (*a*_0_*, b*_0_*, c, a*_1_*, b*_1_) for female (pink) and male (blue) African lions of the Serengeti population. The analysis was conditioned on survival of the first 3 months of life.

**Figure 4:**
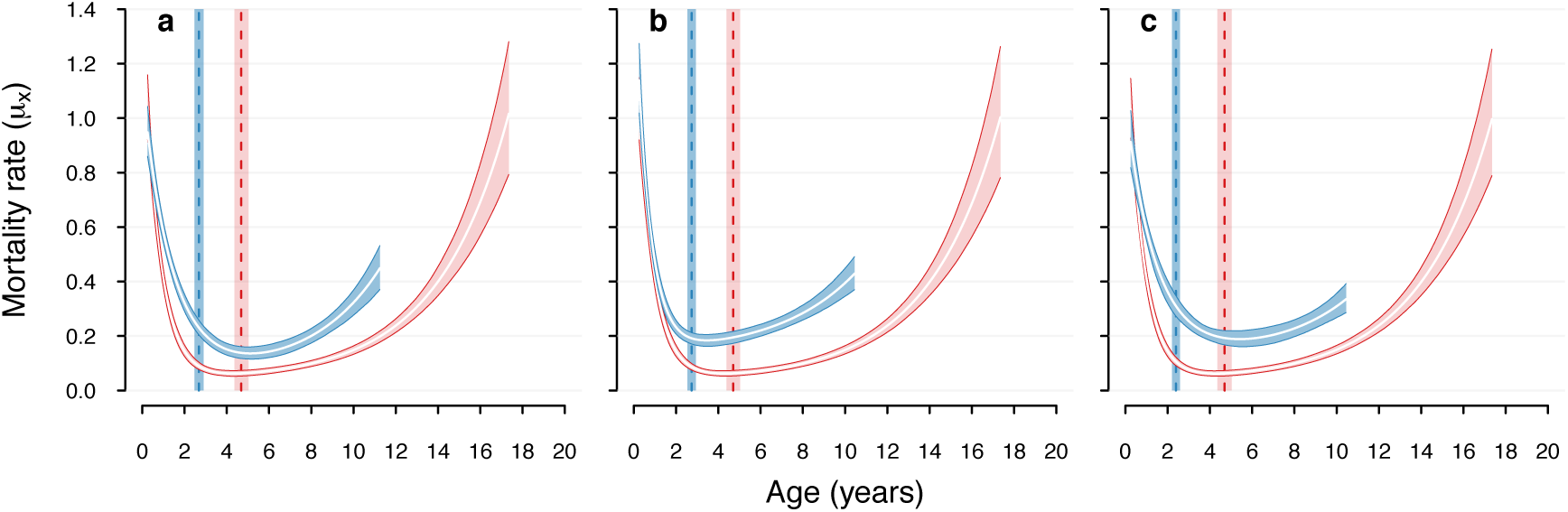
Age-specific mortality estimates for male (blue lines and polygons) and female African lions (pink lines and polygons) of the Serengeti population. Polygons represent 95 % credible intervals of age-specific mortality rates with white lines indicating the mean. Mortality rates are plotted until the ages when 95 % of a synthetic same-sex cohort would be dead. The vertical dashed lines indicate mean life expectancy at 0.25 years of age with the 95 % confident bands indicated by the rectangles. (a) Model A: all males with uncertain fate and old enough for dispersal treated as potential dispersers. (b) Model B: all males indicated by an expert as potential dispersers treated as known dispersers. (c) Model C: all males indicated by an expert as potential dispersers treated as potential dispersers.

Now we turn to the comparison between the models with varying settings for potential dispersers. Model A (Figure 4a) treated the data as if no further information was available on dispersal status of males with uncertain fates (i.e, the default setting of the model). Model B took advantage of expert knowledge on lion behaviour and treated all males that a lion expert believed were dispersers, as known dispersers (Figure 4b). Finally, Model C treated all expert-indicated potential dispersers as potential dispersers (Figure 4c.) The number of potential dispersers whose dispersal state was imputed as a latent state was therefore smaller in Model C when compared to Model A.

We compare these models by examining the estimated mortality rates (Figure 4), the posterior density distributions (Figure 3), and the KL divergences (Figure 5). Since females were treated the same way in all three models, the posterior distributions of parameters for females were congruent among the three models (Figure 3). This was well-captured by the corresponding KL divergences, which were close to, or equal to, 0.5 (Figure 5). Consequently, female mortality rates were almost identical across all three models (Figure 4).

**Figure 5:**
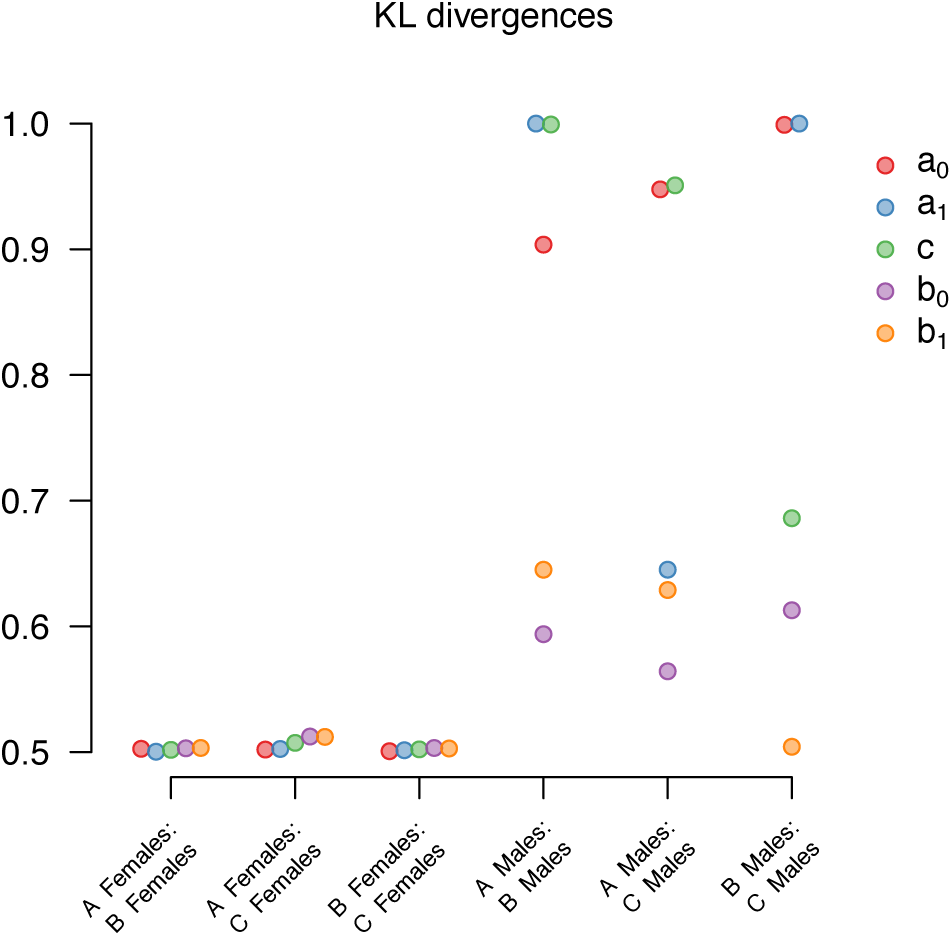
Kullback-Leibler (KL) divergences comparing same-sex Siler parameter posteriors among the three models (A, B, C) with varying settings for males with uncertain fate. Note that the KL divergence estimates are jittered in x-axis direction to improve visibility. The analysis was conditioned on survival of the first three months of life.

For males, the three models gave slightly varying results. The different settings regarding potential dispersers mostly affected the estimation of the Siler parameters that describe initial mortality (*a*_0_), the age-dependent decrease in mortality at young ages (*a*_1_), and the age-independent mortality (*c*) (Figure 5). The initial mortality was higher in Model B, and lower in Model C, when compared to the default model A (Figure 3, Table S1). The age-dependent decrease in mortality was steeper in Model B compared to Model A but similar between Model A and C. The age-independent mortality was higher in both Model B and C when compared to the default Model A. The parameters governing mortality rates at older age (*b*_0_ and *b*_1_) were similar across all three models and the confidence bands overlapped (Figure 3, Table S1).

Because the Siler mortality model is a composite of three additive mortality hazards, the differences among the three models can be more fully understood by comparing the male mortality rates predicted from the three models (Figure 4). Due to the steep decline in age-dependent mortality at younger ages when all expert-indicated dispersers were treated as dispersers (Model B), mortality rates during the juvenile ages up to approximately three years of age were lower in Model B when compared to both models that imputed dispersal state for potential dispersers (Model A and C). However, for the prime-adult-ages, Model B gave the highest mortality estimates, followed by Model C, and then Model A, which gave the lowest estimates. Mortality rates at older ages were highest in Model A and B. Despite these differences in the shape of the mortality rates curves, the life expectancies at 0.25 years of age were predicted to be identical by Model A and B (2.7 years), and only slightly different by Model C (2.4 years).

## Discussion

Life history data of wild animals are often incomplete because animals, even though alive and well, may temporarily or permanently be absent when researchers try to observe them at a given location. This simple observation has far reaching consequences for the estimation of biological properties from these data. Accordingly, various statistical approaches have been developed that account for temporal and spatial heterogeneity in recapture probabilities. For example, multistate CMRR methods have been applied to estimate survival rates while accounting for migration between locations within study sites (Arnason, 1973; Schwarz, Schweigert & Arnason, 1993; Lebreton & Pradel, 2002; Pradel, 2005; Mackenzie *et al*., 2009; Lagrange *et al*., 2014). And spatially explicit CMRR methods have been developed to estimate survival probabilities and population size (Borchers & Efford, 2008; Efford & Mowat, 2014; Ergon & Gardner, 2014). Furthermore, a recently developed spatially explicit Cormack-Jolly-Seber approach jointly models dispersal and survival hierarchically for species in which dispersal movements can be assumed to follow a random walk (Schaub & Royle, 2014).

However, none of these models can account for the extreme case of spatially structured detection probabilities and age-dependent dispersal probabilities that are typical for males of highly-detectable species in which males disperse around the age of maturity. In these species, males are re-sighted with certainty as long as they are alive, and they are not re-sighted after the dispersed. To meet these challenges, our model does not model spatially heterogeneous detection probabilities and dispersal distances but rather imputes the dispersal state of the uncertain male records (i.e., died or dispersed) as a latent state variable in a Bayesian hierarchical framework (Clark *et al*., 2005; Colchero & Clark, 2012; Colchero, Jones & Rebke, 2012). We therefore show that for species with male natal dispersal, mortality and dispersal can be jointly modelled without using movement data. Of course, movement data could potentially be used to inform the dispersal process. However, we decided to develop a model that does not rely on spatial data so that the model can easily be applied to data sets that differ in the structure of available spatial data.

To gauge the possibility of estimating sex- and age-specific mortality in species with male natal dispersal, we focussed on data with incomplete records for sex and age at death. We assumed that this uncertainty could arise form only one of two mechanisms. Firstly, native-born males that disperse from the study area can caused uncertainty in male records of age at death, and secondly, individuals dying as juveniles before their sex could be determined resulted in uncertain sex records. Implicitly, the model therefore assumes that all birth dates are known and that all other types of records can be treated as complete records. Consequently, the model treated the last seen ages of immigrants, native-born females, and males that were imputed to be non-dispersers as certain ages at death. The accuracy of the model therefore hinges on the assumption that only males disperse, and that they disperse only once during their life. During our study, it became apparent that while this assumption holds for some lion populations (A. Loveridge, unpublished data), it does not hold for the Serengeti population. Relaxing the assumption and accounting for higher-order dispersal necessitates a customised extension of the mortality model we present here. The effectiveness of fitting this more complex model depends on the availability of information on both known deaths and dispersal events among immigrants. In the case of the Serengeti population, we took advantage of the expert’s indication on likely dispersal state of disappearing immigrants and extended the default model (Model A) to treat all immigrants that were indicated to be likely dispersers as censored at last seen ages. The difference between the male mortality estimates from the default model and the extended model provides an indication of the amount by which male mortality is overestimated if secondary dispersal is not accounted for (Figure 6).

**Figure 6:**
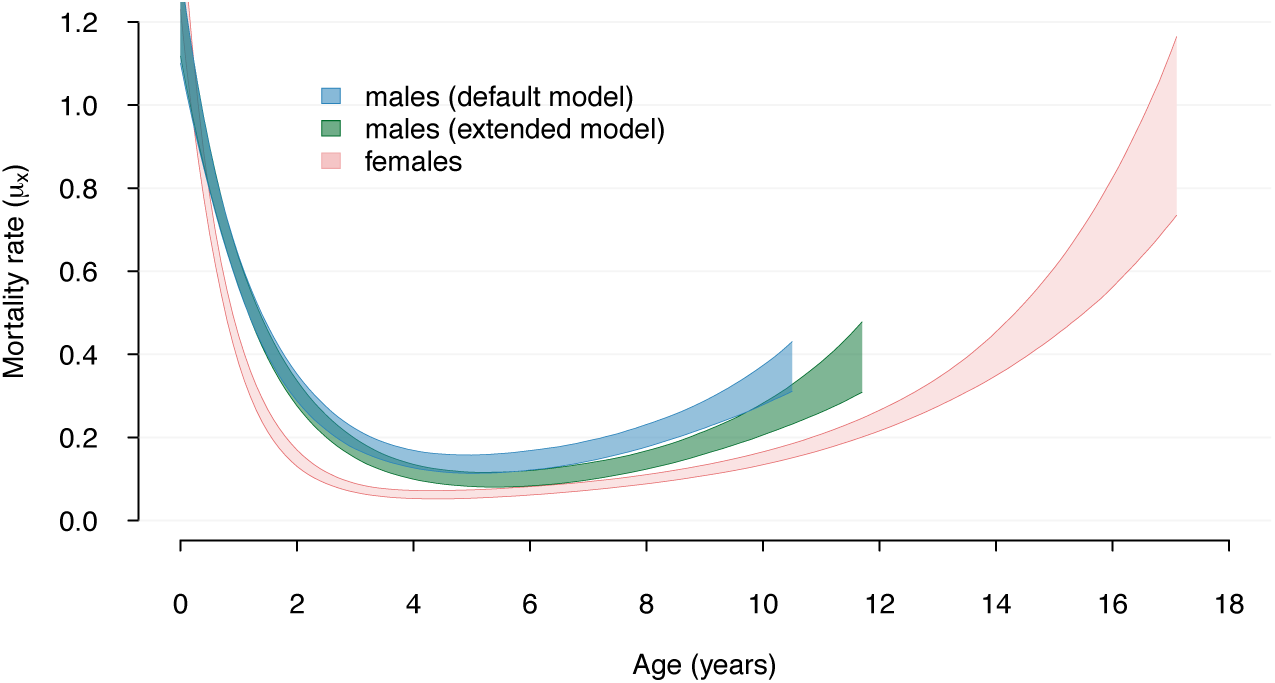
Age-specific mortality estimates for male and female African lions of the Serengeti population. Polygons represent 95 % credible intervals of age-specific mortality rates. Mortality rates are plotted until the ages when 95 % of a synthetic same-sex cohort would be dead. The blue polygons represent male mortality rates obtained from the default model that accounts for natal dispersal. The green polygons represent male mortality rates obtained from an extended model, where secondary dispersal was accounted for additionally to natal dispersal by entering last seen ages of likely secondary dispersers as age of right-censoring.

Another consequence of the treatment of immigrants’ last seen ages as ages at death is that the ratio of immigrants to dispersers is likely to influence the estimation of male mortality parameters. This causes problems if the probability of dispersal out of the study area is much higher than the probability of immigrating into it (see Figure S1 for a simulation). This may be the case for field sites that are established in protected areas and act as a source population for surrounding habitats of lower quality. Mortality in these habitats, and mortality during the dispersal process itself, may also be higher than mortality within the study area. Our model cannot account for this heterogeneity.

Overall, given the recovery of the mortality parameters in the simulation study, we conclude that our approach is a promising step towards obtaining unbiased estimates of male mortality for species with incomplete male records due to male natal dispersal. Mortality parameters were recovered even if sample sizes were low, observed deaths among potential dispersers were rare, and some individuals died before sex could be determined. Furthermore, the comparison of the different models for the lion data allows us to draw some conclusions about the sensitivity of mortality estimates to varying levels of uncertainty in male records. If all expert-indicted dispersers were in fact dispersers (Model B), then by comparing the mortality rates estimated by this model and by the one with the default treatment of dispersers (Model A), we learn that the default model may have the tendency to overestimate mortality during juvenile ages (lower *a*_1_ in Model A than B). The default model may furthermore slightly underestimate mortality during prime-adult ages. Since the model that treats all expert-indicated dispersers as potential dispersers (Model C) shares properties of both Models A and B (similar *c* to Model B, similar *a*_0_ and *a*_1_ to Model A), and may come closest to reality, it seems like a promising avenue for future development to directly include expert knowledge in the Bayesian framework via priors. However, as with the use of spatial data to inform the model, this information is an idiosyncrasy of the data set that we used here. Making the model dependend on this information would therefore preclude the application of the model to estimate mortality for other populations and species. Despite the differences in mortality estimates obtained from the three models, all three models predicted very similar life expectancies, and the confidence bands of predicted mortality rates overlapped across many ages. We therefore conclude that our model provides a good solution to the challenge of estimating male mortality in species with data-deficiency for males due to natal dispersal.

## Acknowledgements

JAB acknowledges funding from the International Max Planck Research Network on Aging (MaxNetAging). JAB thanks Owen Jones, Jacques Deere, Emily Simmonds, and Tim Coulson for helpful comments on the manuscript.

## Supporting Information

Methods S1 Calculation and calibration of Kullback-Leibler divergence

Figure S1 Traces of mortality and dispersal parameter estimation for Models A to C.

Figure S2 Predicted mortality rates for simulated data if male immigration probability was set to 0.5.

Table S1 Estimated coefficients for models A to C.

Code S1 R code to simulate data. Download from github.com/bartholdja/DPhil_supplements.

Code S2 R code to run the model on simulated data. Download from github.com/bartholdja/DPhil_supplements.

## Supporting methods, figures, and tables

### Methods S1: Calculation and calibration of Kullback-Leibler divergence

The Kullback-Leibler (KL) divergence calculates the difference or the amount of overlap between two distributions (Kullback & Leibler, 1951; McCulloch, 1989; Burnham & Anderson, 2001). To illustrate the calculation of KL, let’s take a parameter *θ*, for which the resulting ‘sub-parameters’ for females and males would be *θ_f_* and *θ_m_*, respectively. Thus, for an individual *i*, we have *θ* = *θ_f_I_i_* + θ_m_(1 − *I_i_*), where *I_i_* is an indicator function that assigns 1 if the individual is a female and 0 otherwise. For each of these parameters, our model produces a posterior distribution, say *P_f_* = *p*(*θ_f_*|…) and *P_m_* = *p*(*θ_m_*|…), respectively. The KL between these distributions is calculated as

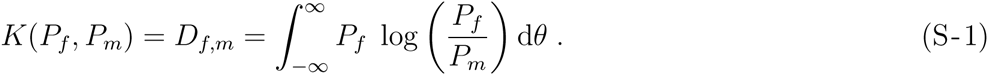

The result can be interpreted as how far off we would be if we tried to predict *θ_m_* one from the posterior distribution of *θ_f_*. If both distributions are identical, then *D_f_,_m_* = 0, suggesting that there is no distinction between the parameters of both covariates and hence, that both covariates have the same effect. With increasing KL values, the discrepancy becomes higher. As can be inferred from Equation S-1, the relationship is asymmetric, namely *K*(*P_f_, P_m_*) ≠ *K*(*P_m_,P_f_*).

To make KL values easier to interpret, McCulloch (1989) proposed a simple calibration of the KL values that reduces the asymmetry. Let *k* = *K*(*P_f_,P_m_*) and *q_k_* be a calibration function such that

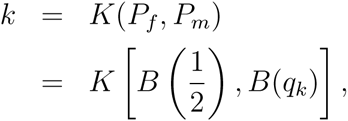
where 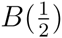 is a Bernouilli distribution for an event with probability 0.5 (i.e., same probability of success and failure). This calibration is then calculated as

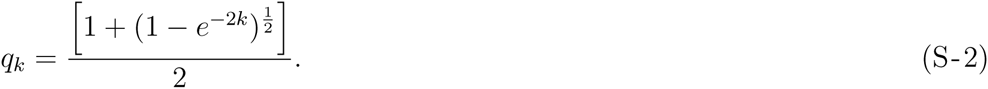

Thus, *q_k_* ranges from 0.5 to 1, where a value of 0.5 means that the distributions are identical, and 1 that there is no overlap between them.

**Figure S1:**
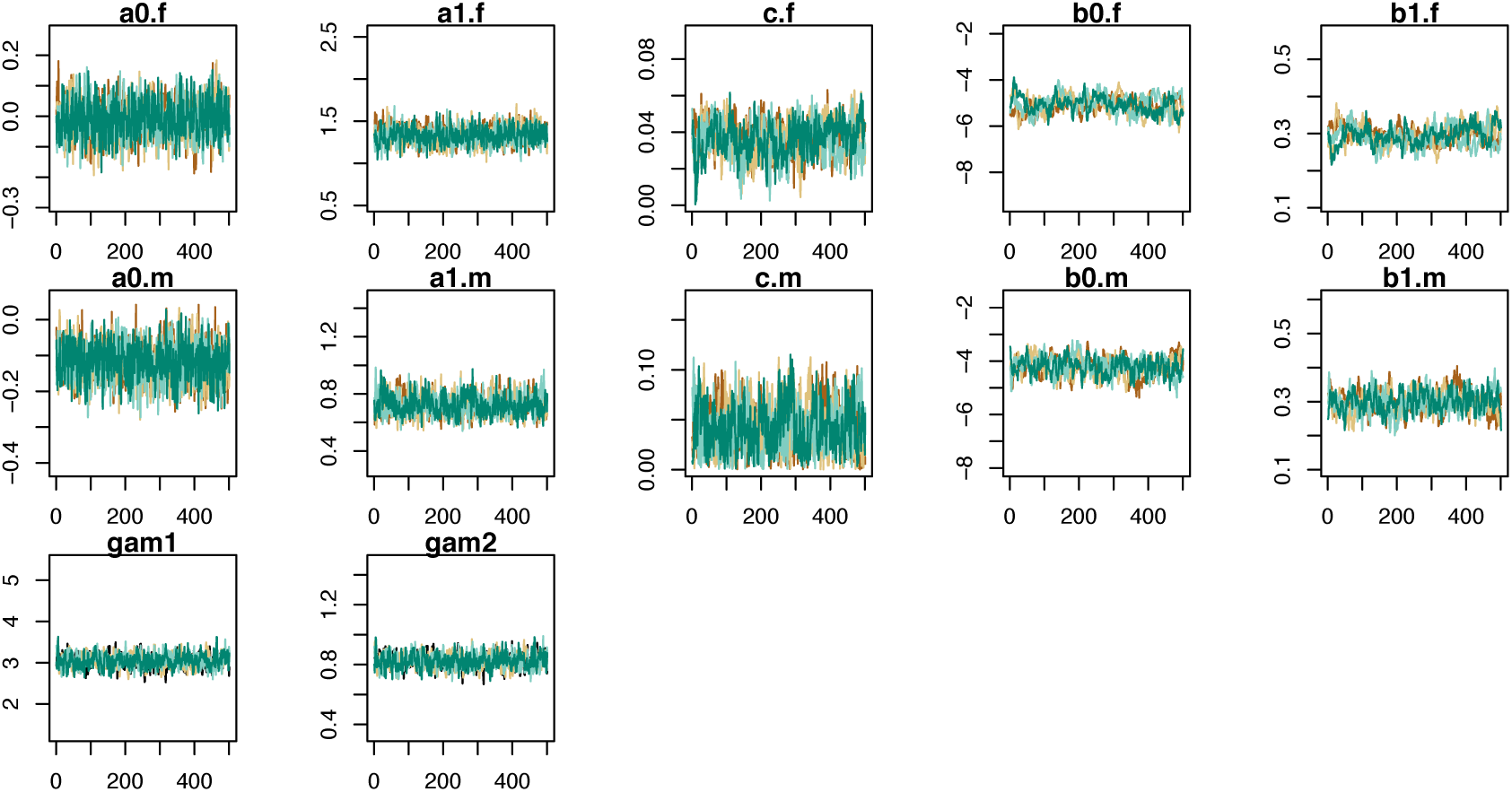
Trace plots for four parallel runs for the Serengeti lion mortality analysis. Estimated parameters are the Siler parameters (*a*_0_, *b_0_, c, a*_1_*, b*_1_; f denotes estimates for females and m for males) and Gamma parameters (shape and rate; gam1 and gam2). Model A: all males with uncertain fate and last seen ages older than minimum age at dispersal treated as potential dispersers.

**Figure S2:**
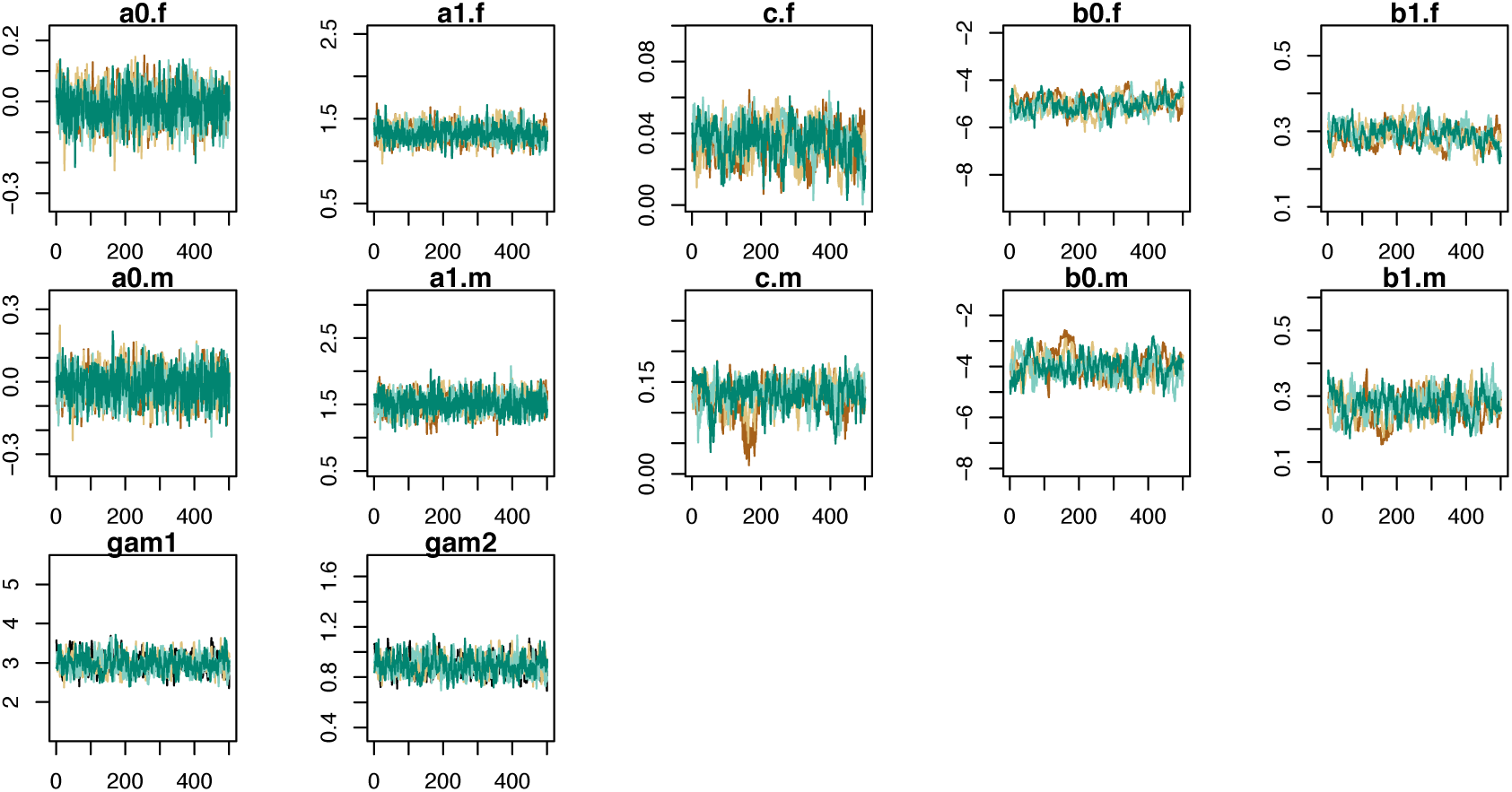
Trace plots for four parallel runs for the Serengeti lion mortality analysis. Estimated parameters are the Siler parameters (*a*_0_*, b*_0_*, c, a*_1_*, b*_1_; f denotes estimates for females and m for males) and Gamma parameters (shape and rate; gam1 and gam2). Model B: all males that an expert indicated as potential dispersers treated as known dispersers.

**Figure S3:**
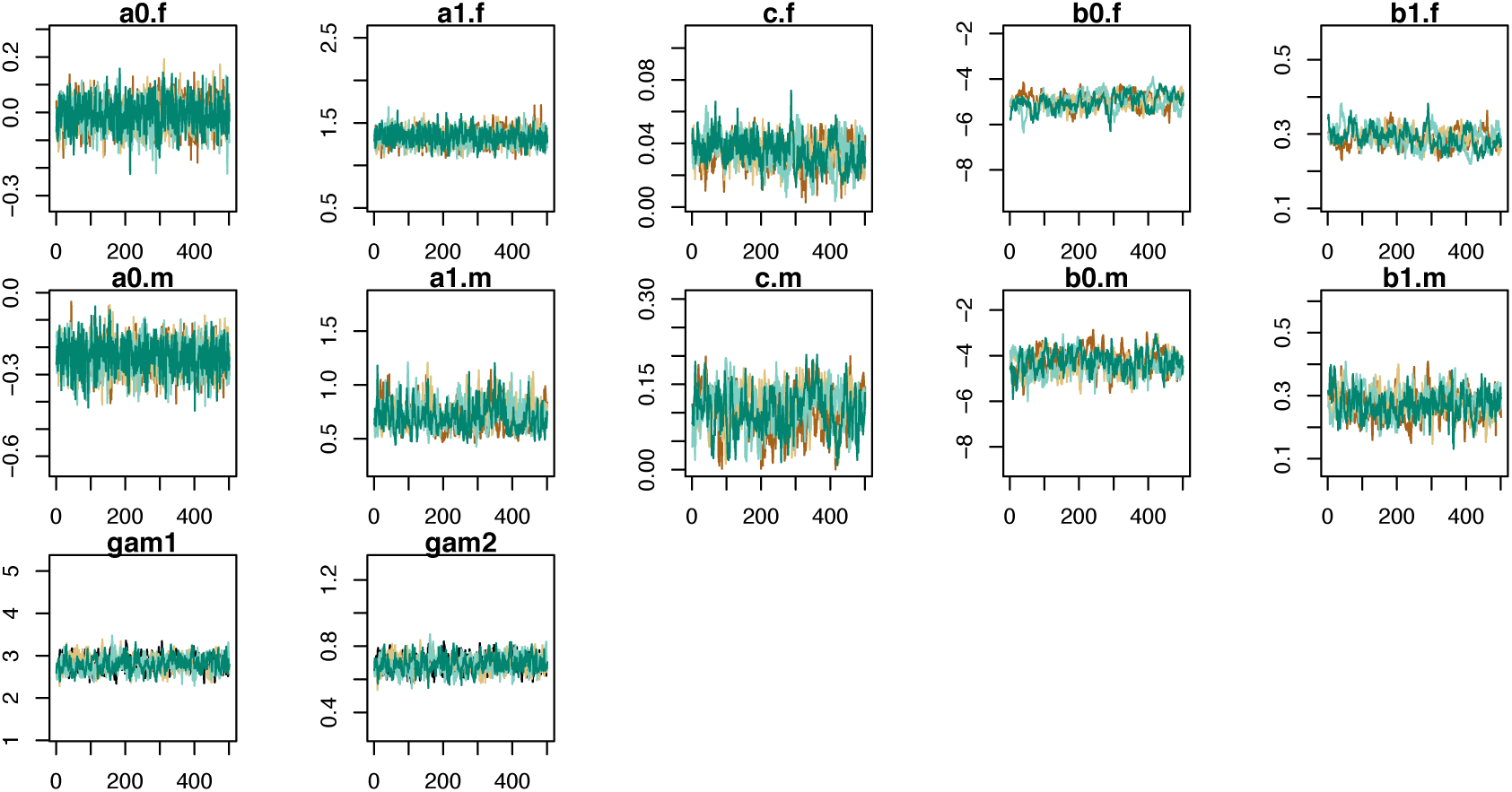
Trace plots for four parallel runs for the Serengeti lion mortality analysis. Estimated parameters are the Siler parameters (*a*_0_*, b*_0_*, c, a*_1_*, b*_1_; f denotes estimates for females and m for males) and Gamma parameters (shape and rate; gam1 and gam2). Model C: All males that an expert indicated as potential dispersers treated as potential dispersers.

**Figure S4:**
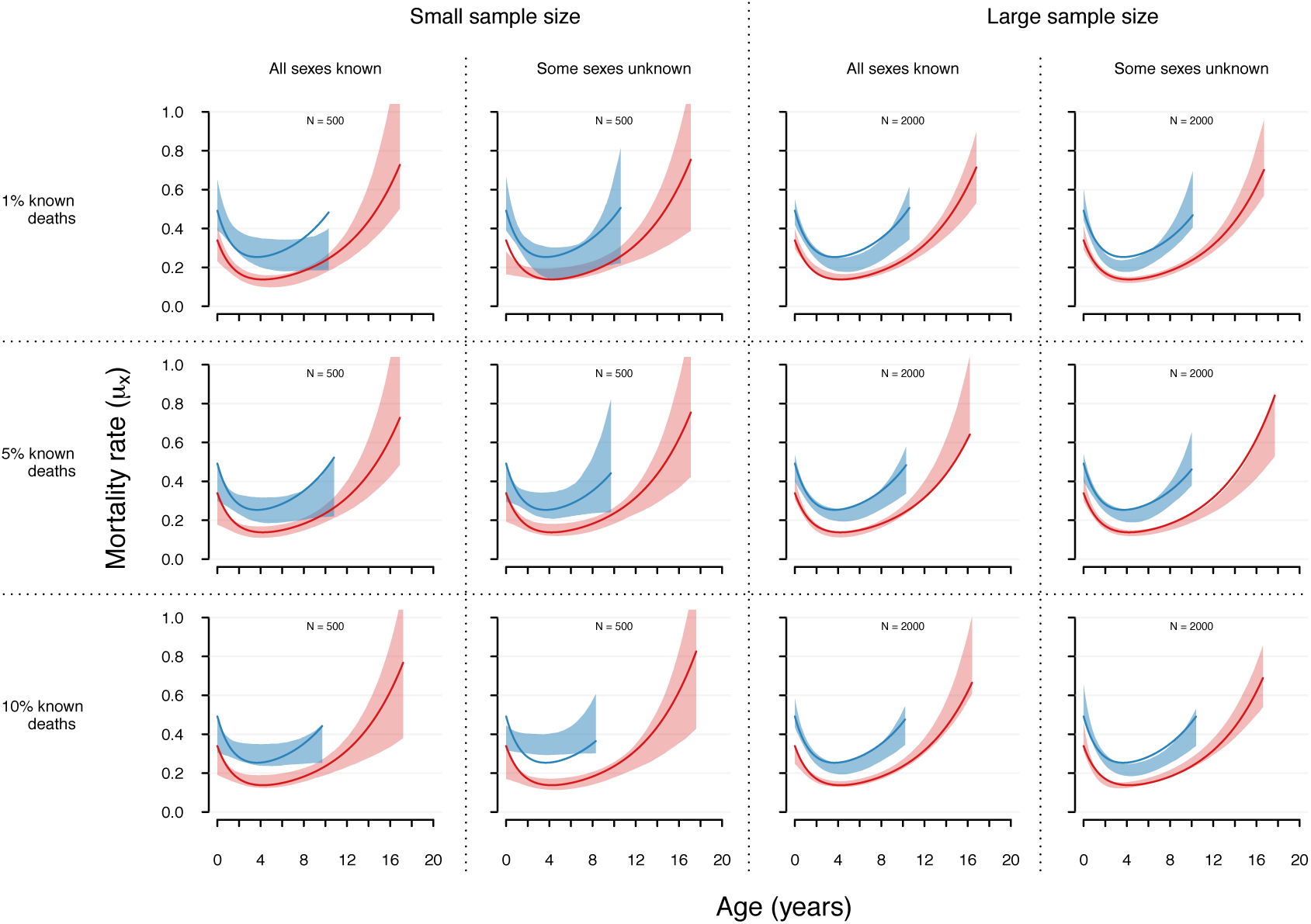
Predicted mortality functions for males (blue polygons) and females (pink polygons) compared to the mortality functions used to simulate the data (solid lines), if the probability to immigrate into the study area of males born outside of it was lowered from 1 to 0.5. Polygons represent 95 % credible intervals of age-specific mortality functions. Mortality rates are plotted until the ages when 95 % of a synthetic same-sex cohort would be dead. Results are given for 12 simulations varying the size of the native-born population (N = 500 or N = 2000), the proportion of known deaths among last seen ages (1%, 5%, or 10%), and whether the sex of 30% of individuals dying younger than 1 year of age remained undetermined or not.

**Table S1:** Estimated Siler and gamma coefficients for Model A (all males with uncertain fate and old enough for dispersal treated as potential dispersers), Model B (all males indicated by an expert as potential dispersers treated as known dispersers, and Model C (all males indicated by an expert as potential dispersers treated as potential dispersers). Given are mean, SE, and credible intervals of the parameter posterior distributions.

|  |  | Coefficient | Mean | SE | 2.5 % | 97.5 % |
| --- | --- | --- | --- | --- | --- | --- |
| Model A | Females | $a_0$ | -0.01 | 0.06 | -0.12 | 0.11 |
| | | $a_1$ | 1.33 | 0.10 | 1.14 | 1.54 |
| | | $c$ | 0.04 | 0.01 | 0.02 | 0.05 |
| | | $b_0$ | -5.06 | 0.36 | -5.79 | -4.38 |
| | | $b_1$ | 0.29 | 0.03 | 0.24 | 0.35 |
| | Males | $a_0$ | -0.12 | 0.05 | -0.22 | -0.01 |
| | | $a_1$ | 0.72 | 0.07 | 0.60 | 0.87 |
| | | $c$ | 0.04 | 0.02 | 0.00 | 0.09 |
| | | $b_0$ | -4.19 | 0.35 | -4.89 | -3.55 |
| | | $b_1$ | 0.30 | 0.03 | 0.24 | 0.36 |
| | | $gam_1$ | 3.04 | 0.17 | 2.72 | 3.37 |
| | | $gam_2$ | 0.82 | 0.05 | 0.73 | 0.92 |
| Model B | Females | $a_0$ | -0.01 | 0.06 | -0.12 | 0.10 |
| | | $a_1$ | 1.33 | 0.10 | 1.14 | 1.53 |
| | | $c$ | 0.04 | 0.01 | 0.01 | 0.05 |
| | | $b_0$ | -5.02 | 0.36 | -5.70 | -4.33 |
| | | $b_1$ | 0.29 | 0.03 | 0.24 | 0.34 |
| | Males | $a_0$ | -0.01 | 0.07 | -0.14 | 0.12 |
| | | $a_1$ | 1.52 | 0.14 | 1.25 | 1.80 |
| | | $c$ | 0.13 | 0.02 | 0.07 | 0.17 |
| | | $b_0$ | -3.98 | 0.44 | -4.81 | -3.03 |
| | | $b_1$ | 0.27 | 0.04 | 0.19 | 0.34 |
| | | $gam_1$ | 2.97 | 0.22 | 2.56 | 3.45 |
| | | $gam_2$ | 0.89 | 0.07 | 0.76 | 1.04 |
| Model C | Females | $a_0$ | -0.01 | 0.06 | -0.12 | 0.10 |
| | | $a_1$ | 1.33 | 0.09 | 1.15 | 1.51 |
| | | $c$ | 0.03 | 0.01 | 0.02 | 0.05 |
| | | $b_0$ | -4.99 | 0.34 | -5.68 | -4.35 |
| | | $b_1$ | 0.29 | 0.03 | 0.24 | 0.34 |
| | Males | $a_0$ | -0.24 | 0.06 | -0.36 | -0.12 |
| | | $a_1$ | 0.71 | 0.12 | 0.51 | 1.00 |
| | | $c$ | 0.11 | 0.04 | 0.03 | 0.17 |
| | | $b_0$ | -4.31 | 0.47 | -5.21 | -3.34 |
| | | $b_1$ | 0.27 | 0.04 | 0.19 | 0.36 |
| | | $gam_1$ | 2.81 | 0.18 | 2.46 | 3.16 |
| | | $gam_2$ | 0.69 | 0.05 | 0.60 | 0.80 |

## References

Andersson, M. (1994) Sexual selection. Princeton University Press, Princeton (NJ).

Arnason, A.N. (1973) The estimation of population size, migration rates and survival in a stratified population. Researches on Population Ecology, 15, 1–8.

Borchers, D.L. & Efford, M.G. (2008) Spatially explicit maximum likelihood methods for capture-recapture studies. Biometrics, 64, 377–385.

Burnham, K.P. & Anderson, D.R. (2001) Kullback-Leibler information as a basis for strong inference in ecological studies. Wildlife Research, 28, 111–119.

Clark, J.S., Ferraz, G.A., Oguge, N., Hays, H. & DiCostanzo, J. (2005) Hierarchical Bayes for structured, variable populations: from recapture data to life-history prediction. Ecology, 86, 2232–2244.

Clark, J.S. (2007) Models for ecological data. Princeton University Press, Princeton (NJ).

Colchero, F. & Clark, J.S. (2012) Bayesian inference on age-specific survival for censored and truncated data. Journal of Animal Ecology, 81, 139–149.

Colchero, F., Jones, O.R. & Rebke, M. (2012) BaSTA an R package for Bayesian estimation of age-specific survival from incomplete mark-recapture/recovery data with covariates. Methods in Ecology and Evolution, 3, 466–470.

Cormack, R.M. (1964) Estimates of survival from the sighting of marked animals. Biometrika, 51, 429–438.

Cubaynes, S., Pradel, R., Choquet, R., Duchamp, C., Gaillard, J.M., Lebreton, J.D., Marboutin, E., Miquel, C., Reboulet, A.M., Poillot, C., Taberlet, P. & Gimenez, O. (2010) Importance of accounting for detection heterogeneity when estimating abundance: the case of French wolves. Conservation Biology, 24, 621–626.

Dupuis, J.A. (2002) Prior distributions for stratified capture-recapture models. Journal of Applied Statistics, 29, 225–237.

Dupuis, J.A. (1995) Bayesian estimation of movement and survival probabilities from capture-recapture data. Biometrika, 82, 761–772.

Efford, M.G. & Mowat, G. (2014) Compensatory heterogeneity in spatially explicit capture-recapture data. Ecology, 95, 1341–1348.

Ergon, T. & Gardner, B. (2014) Separating mortality and emigration: modelling space use, dispersal and survival with robust-design spatial capture-recapture data. Methods in Ecology and Evolution, 5, 1327–1336.

Gelfand, A.E. & Smith, A.F.M. (1990) Sampling-based approaches to calculating marginal densities. Journal of the American Statistical Association, 85, 398–409.

Jolly, G.M. (1965) Explicit estimates from capture-recapture data with both death and immigration-stochastic model. Biometrika, 52, 225–247.

King, R. & Brooks, S.P. (2002) Bayesian model discrimination for multiple strata capture-recapture data. Biometrika, 89, 785–806.

Kullback, S. & Leibler, R.A. (1951) On information and sufficiency. Annals of Mathematical Statistics, 22, 142–143.

Lagrange, P., Pradel, R., Belisle, M. & Gimenez, O. (2014) Estimating dispersal among numerous sites using capture-recapture data. Ecology, 95, 2316–2323.

Le Galliard, J.F., Fitze, P.S., Ferriere, R. & Clobert, J. (2005) Sex ratio bias, male aggression, and population collapse in lizards. Proceedings of the National academy of Sciences, 102, 18231–18236.

Lebreton, J. & Pradel, R. (2002) Multistate recapture models: modelling incomplete individual histories. Journal of Applied Statistics, 29, 353–369.

Mackenzie, D.I., Nichols, J.D., Seamans, M.E. & Gutierrez, R.J. (2009) Modeling species occurrence dynamics with multiple states and imperfect detection. Ecology, 90, 823–835.

McCulloch, R.E. (1989) Local model influence. Journal of the American Statistical Association, 84, 473–478.

Milner-Gulland, E.J., Bukreeva, O.M., Coulson, T., Lushchekina, A.A., Kholodova, M.V., Bekenov, A.B. & Grachev, I.A. (2003) Conservation: reproductive collapse in saiga antelope harems. Nature, 422, 135–135.

Mosser, A., Fryxell, J.M., Eberly, L. & Packer, C. (2009) Serengeti real estate: density vs. fitness-based indicators of lion habitat quality. Ecology Letters, 12, 1050–1060.

Mysterud, A., Coulson, T. & Stenseth, N.C. (2002) The role of males in the dynamics of ungulate populations. Journal of Animal Ecology, 71, 907–915.

Packer, C. (2005) Ecological change, group territoriality, and population dynamics in Serengeti lions. Science, 307, 390–393.

Packer, C., Pusey, A.E., Rowley, H., Gilbert, D.A., Martenson, J. & O’Brien, S.J. (1991) Case-study of a population bottleneck - Lions of the Ngorongoro Crater. Conservation Biology, 5, 219–230.

Pradel, R. (2005) Multievent: An extension of multistate capture-recapture models to uncertain states. Biometrics, 61, 442–447.

Rankin, D.J. & Kokko, H. (2007) Do males matter? The role of males in population dynamics. Oikos, 116, 335–348.

Schaub, M. & Royle, J.A. (2014) Estimating true instead of apparent survival using spatial Cormack-Jolly-Seber models. Methods in Ecology and Evolution, 5, 1316–1326.

Schwarz, C.J., Schweigert, J.F. & Arnason, A.N. (1993) Estimating migration rates using tag-recovery data. Biometrics, 49, 177–193.

Seber, G.A. (1965) A note on the multiple-recapture census. Biometrika, 52, 249–259.

Siler, W. (1979) A competing-risk model for animal mortality. Ecology, 60, 750–757.

Smuts, G.L., Anderson, J.L. & Austin, J.C. (1978) Age determination of the African lion (*Panthera leo*). Journal of Zoology, 185, 115–146.

White, G.C. & Burnham, K.P. (1999) Program MARK: survival estimation from populations of marked animals. Bird Study, 46, 120–139.

Whitman, K., Starfield, A.M., Quadling, H.S. & Packer, C. (2004) Sustainable trophy hunting of African lions. Nature, 428, 175–178.

